# Redefining phenotypic intratumor heterogeneity of pancreatic ductal adenocarcinoma: a bottom-up approach

**DOI:** 10.1101/2023.11.16.567454

**Authors:** Marc Hilmi, Flore Delecourt, Jérôme Raffenne, Taib Bourega, Nelson Dusetti, Juan Iovanna, Yuna Blum, Magali Richard, Cindy Neuzillet, Anne Couvelard, Louis de Mestier, Vinciane Rebours, Rémy Nicolle, Jérôme Cros

## Abstract

**Background:** Pancreatic ductal adenocarcinoma (PDAC) tumor inter-patient heterogeneity has been well described with two major prognostic subtypes (classical and basal-like). An important intra-patient heterogeneity has been reported but has not yet been extensively studied due to the lack of standardized, reproducible and easily accessible high throughput methods.

**Material and Methods:** We built an immunohistochemical (IHC) tool capable of differentiating RNA-defined classical and basal-like tumors by selecting relevant antibodies using a multi-step process. The successive stages of i) an *in-silico* selection from a review literature and a bulk transcriptome analysis of 309 PDACs, ii) a tumor-specific selection from 30 patient-derived xenografts followed by iii) the validation on tissue microarrays in 50 PDAC were conducted. We used our final IHC panel on two independent cohorts of resected PDAC (n=95, whole-slide, n=148, tissue microarrays) for external validation. After digitization and registration of pathology slides, we performed a tile-based-analysis in tumor and pre-neoplastic epithelial areas and a k-means clustering to identify relevant marker combinations.

**Results:** Sequential marker selection led to the following panel: GATA6, CLDN18, TFF1, MUC16, S100A2, KRT17, PanBasal. Four different phenotypes were identified: 1 classical, 1 intermediate (KRT17+) and 2 basal-like (MUC16+ vs S100A2+) with specific biological properties. The presence of a minor basal contingent drastically reduced overall survival, even in classical predominant PDACs (HR=2.36, p=0.01). Analysis of preneoplastic lesions suggested that pancreatic carcinogenesis may follow a progressive evolution from classical toward a basal through an early intermediate phenotype.

**Conclusion:** Our IHC panel redefined and easily assessed the high degree of intra- and inter-tumoral heterogeneity of PDAC.

## Introduction

Pancreatic ductal adenocarcinoma (PDAC) has one of the poorest prognosis, due to its diagnosis often made at a late stage and low sensitivity to therapies. Two major PDAC transcriptomic subtypes have been described: a classical subtype, which expresses epithelial differentiation genes, with a better prognosis than the basal-like subtype, expressing epithelial-mesenchymal transition genes(1). Yet, recent studies questioned the existence of such a blunt dichotomy at the tumor cell level by highlighting frequently misclassified or hybrid/mixed tumors (2–4).

PDAC displays not only inter-patient but also intra-patient heterogeneity. At the DNA level, different clonal populations harboring specific mutations were observed in PDAC(5). From an histological standpoint, PDAC is heterogenous as different histological architectures have been described and can coexist within the same tumor introducing the concept of intra-tumoral epithelial heterogeneity(6, 7). The squamous/non-gland forming morphology is associated with the basal-like subtype, whereas the glandular/gland-forming morphology is associated with the classical subtype(7, 8). However, within the classical transcriptomic subgroup, there seem to be tumors or parts of tumors that do not form glands and have a worse prognosis than purely glandular tumors(2, 7). At the RNA level, bulk transcriptomic analyses are costly, difficult to implement in clinical practice, and gives a global reflection of the tumor, without being able to reliably highlight any heterogeneity. Consequently, a continuous molecular gradient was proposed to better recapitulate the histological groups, allowing to order the tumors based on their “classicness or basalness” but with no information on intra-tumor heterogeneity(9). Several single cell studies helped to decipher intratumor heterogeneity and suggested that there is a co-occurrence of the classical and basal-like subtypes within the same PDAC (hybrid or mixed tumors) (3, 8, 10, 11). Moreover, Williams et al.(11) showed that cells co-expressing classical and basal markers are present in over 90% of PDACs and could constitute an intermediate phenotype. However, these approaches have been applied to a small number of patients restricting their conclusions. Our team has recently showed the possibility of classifying cases into subtypes based on histology features from the hematoxylin and eosin stain (H&E) through a deep learning model(2). In addition to the hybrid tumors combining classical and basal components, intermediate tumors for which most tumor cells could not be morphologically clearly assigned to any of the two classical and basal subtypes were revealed. Nevertheless, the morphological approach is time consuming to fully assess the extent of intratumor heterogeneity and requires expertise in pancreatic pathology. Multi areas microdissections are an interesting approach for exploring intratumoral heterogeneity but are unable to evaluate heterogeneity within a single gland. Overall, the definition and estimation of intra-tumoral heterogeneity on a large scale is currently not feasible due to the lack of a standardized and an easily accessible method.

The strength of immunohistochemistry (IHC), compared to transcriptomic or HES analysis of tumors, lies in the fact that this technique allows, in an easy and cost-effective way, a fine analysis of each tumor cell conserving its spatial localization. Our objective was to build a robust IHC panel reflecting molecular subtypes in order to explore PDAC intratumoral heterogeneity.

## Material and methods

### Patient samples

The *in-silico* marker selection was performed on a retrospective multicenter cohort including 381 resected PDAC in four university hospitals (Saint-Antoine (APHP), La Pitié Salpêtrière (APHP), Ambroise Paré (APHP), and Erasmus (Brussels)) between September 1996 and December 2010 (12). Briefly, exclusion criteria were neoadjuvant chemotherapy or radiochemotherapy, macroscopically incomplete surgical tumor resection (R2), and a histologic diagnosis other than adenocarcinoma. Patients who died postoperatively or had complications within 30 days after surgery were also excluded. A total of 312 patients were included in our study. The analysis was subsequently restricted to 309 patients after removing three tumors with unsatisfactory quality controls. The bulk transcriptome of these tumors was obtained by microarrays (Affymetrix HGU219) as previously described(12). From this cohort, 13 clear classical and 8 clear basal-like tumors from the transcriptomic point of view were selected to construct a tissue microarray (TMA) with 4 spots of 0.6mm/case.

The previously published transcriptomes of thirty patient-derived xenografts, for which the human tumor compartment was separated from the mouse stromal compartment and tumor subtyping was performed on the tumor compartment, was obtained from the PaCaOmics Clinical Trial (NCT01692873)(13).

The human validation of the IHC markers on whole slides was performed on a cohort of 50 PDAC cases from Beaujon Hospital with the same inclusion criteria as the cohort described above. The bulk transcriptome of these cases was obtained after macrodissection of the whole tumor area on the same block as the one used for the IHC and obtained from FFPE sections using a high-purity FFPE RNA isolation kit (Roche, Basel, Switzerland) following the manufacturer’s protocol. Library preparation was performed using QuantSeq 3’ mRNA-Seq REV (Lexogen GmbH, Vienna, Austria) with an input of 150 ng of total FFPE RNA. To assess the intratumor heterogeneity on a greater scale, the final IHC panel was performed on an additional cohort of 45 cases with the same inclusion criteria leading to a total cohort of 95 patients for the tile-based analysis.

Finally, a cohort of 148 resected PADC already described was used to construct TMA 4 spots of 0.6mm/case) and validate the prognostic impact of the IHC panel(2). The techniques performed on the 3 cohorts of patients with resected PDAC and their aims are presented in the Supplementary Figure 1.

### Immunohistochemistry

All the IHC stainings were performed on a routine automate (Ventana Benchmark ultra, Roche diagnostic, Tuscon AZ, USA). The details of each antibody are listed in the Supplementary Table 1. For the spatial analysis, IHC stainings were performed on 5 serial slides: 3 slides with a duplex IHC to limit the number of slides (GATA6/S100A2 - PanB/TTF1 - KRT17/CLDN18) and 2 slides with a single marker MUC16 and Pancytokeratin. The stainings were quantified using the H-score, defined as the staining intensity (0=null, 1=weak, 2=moderate, 3=strong) multiply by the percentage of positive tumor cells (ranging from 0 to 100%) leading to a score ranging from 0 to 300. A spot without tumor cells was not scored.

### Image analysis

Registration of digitized whole-slide images of serial histology sections was performed using the VALIS software(14). Manual annotation on HES images by an expert pathologist (JC), virtual tile cutting, and pixel detection were conducted using QuPath version 0.4.3(15). The algorithm for pixel detection was based on a pixel classifier and was trained on representative pictures. Filtration of tiles was based on their size and content of PANCK and performed using R software version 4.3.0.

### Statistical analysis

Tile clustering was performed on logged values by the method of k-means using the first components of the principal component analysis counting for more than 5% of the variability. The optimal number of clusters was based on the silhouette method. Clusters expression was classified according to their presence (>1%, as below can be related to detection artifacts) and predominance (>50%, as the major cluster). Co-occurrence of each cluster was assessed using a Fisher test. We used the Kaplan–Meier method to provide median survival times and to graphically demonstrate survival curves. The log-rank test was used to evaluate statistical significance. We used Cox proportional hazards regression to evaluate associations between tumor clusters and overall survival (OS). Potential prognostic factors included in the models were age > 65 years, resection margins, vascular emboli, lymph node, vascular and perineural invasion. Associations among categorical variables were tested using the chi-squared test for large samples (n > 60) and Fisher exact test for small samples (n < 60). Correlations between continuous variables were analyzed using the Spearman test. Pathway analyses were performed with gene set enrichment analysis (GSEA) using the DESeq2 package. For each test, statistical significance was set at a two-sided p value of <0.05. All statistical analyses were performed with R software version 4.3.0.

## Results

### Development of an immunohistochemistry panel to distinguish basal-like and classical subtypes

In order to build a robust IHC tool capable of differentiating classical and basal-like tumor cells, we selected the relevant markers using a multi-step process (Figure 1A). IHC markers should fulfil the following criteria: (i) to be highly correlated with the basal-like or classical subtype, (ii) to be expressed specifically in the corresponding molecular subtype, (iii) to be expressed only by tumor cells and not by stromal cells, and (iv) to have available antibodies corresponding to the encoded protein that produce a staining compatible with a routine use (intensity, ease of interpretation, robustness).

**Figure 1.**
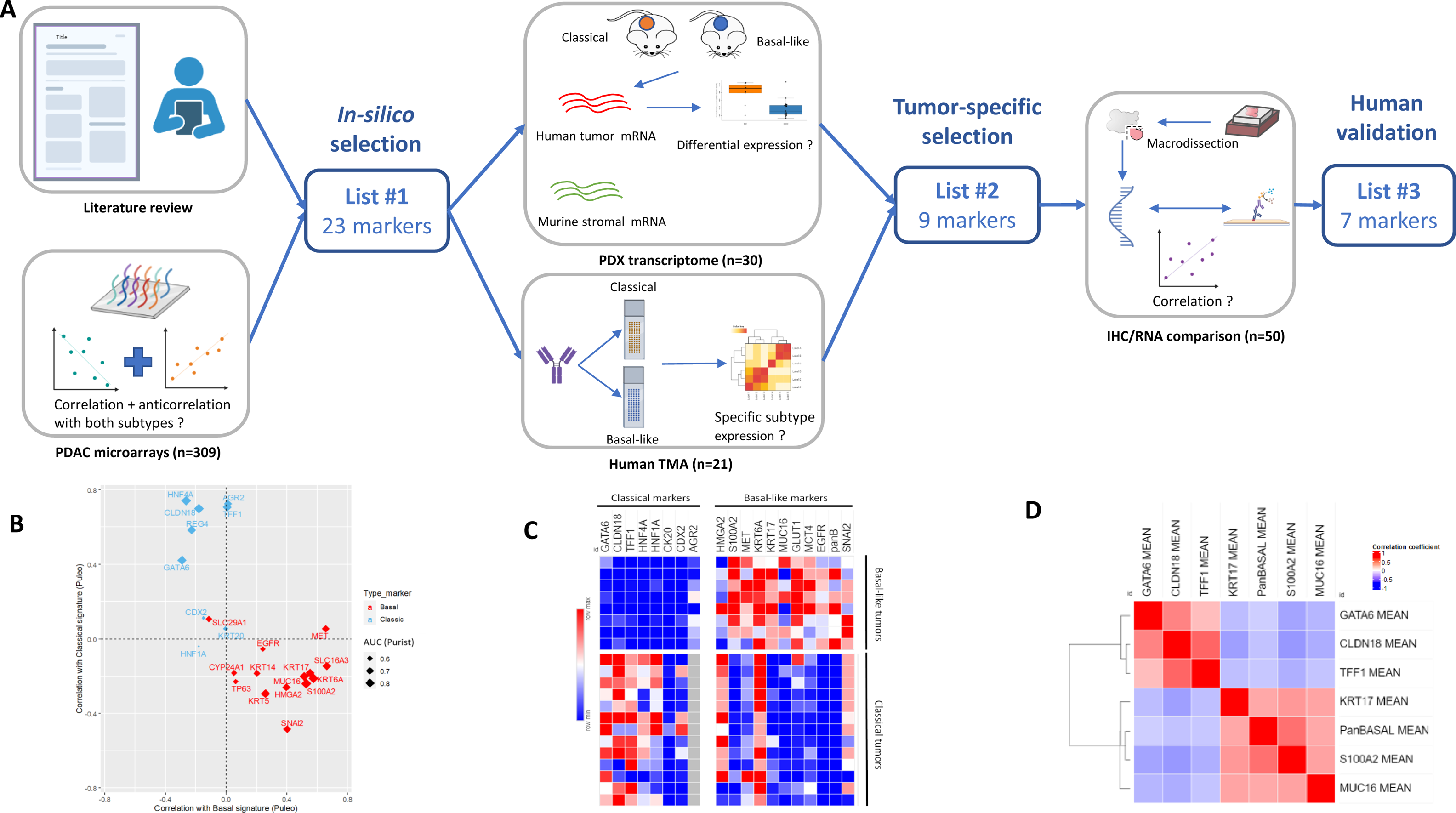
Markers selection to determine the molecular subtype. A. Multi-step process to identify immunohistochemistry markers surrogate of classical and basal-like subtypes. IHC: immunohistochemistry; PDAC: pancreatic ductal adenocarcinoma; PDX: patient-derived xenografts; TMA: tissue microarray. B. Scatter plot showing correlation of selected genes according to the classical and basal signatures from Puleo et al. Size of points is modulated by the value of AUC predicting the Purist signature. C. Expression of tested markers in a set of 13 classical and 8 basal-like pancreatic ductal adenocarcinomas. H-score: 0 (blue) to 300 (red). D. Correlation matrix of the 7 validated immunohistochemistry markers in 50 human pancreatic ductal adenocarcinomas (red = strong correlation, blue = strong anti-correlation).

The first step was to perform an *in-silico* marker selection. To achieve this selection, we performed a literature review to identify surrogate markers of classical and basal subtypes. References for this review were identified through searches of PubMed with the terms “pancreatic cancer” and “molecular subtypes”, from 1995 until September 2023. Only papers published in English were reviewed. Articles were also identified through searches of the authors’ own files. At the end, we identified 23 markers including 9 for the classical subtype (GATA6, TFF1, CLDN18, HNF4A, HNF1A, REG4A, AGR2, KRT20 and CDX2) and 14 for the basal subtype (MET, KRT5, KRT6A, KRT14, S100A2, TP63, SNAI2, KRT17, EGFR, SLC29A1, SLC16A3, MUC16, CYP24A1 and HMGA2) (3, 16–24). In addition, we analyzed transcriptomic data from a retrospectively well-characterized multicenter cohort of 309 patients to explore their correlation and anticorrelation with both subtypes (Figure 1B). Some markers had a low correlation with both subtypes including EGFR, SLC29A1, CYP24A1 and HNF1A.

The second step was to ensure the tumor-specific expression of the 23 markers to support their relevance for subtype distinction. Their corresponding antibodies were assessed on a TMA of 8 and 13 cases selected from the above cohort with homogeneous and strong basal-like and classical signature, respectively. For more convenience, we used one cocktail that combined three markers (KRT5, KRT14 and p63), called PanBasal (PanBS). Representative examples of stainings are presented in the Supplementary Figure 2. The REG4 antibody strongly stained tumor-associated neutrophils and the CYP24A1 antibody gave weak or no staining leading to the exclusion of these markers. The IHC expression of each candidate marker according to basal and classical signatures is shown in Figure 1C. We removed 2 classical (CDX2, KRT20) and 2 basal-like markers (HMGA2, EGFR) because of their weak expression in the corresponding subtype in TMA. We also excluded 6 markers that showed expression in both tumor subtypes (AGR2, SNAI2, KRT6A, MET, SLC29A1, SLC16A3). Finally, 5 classical markers (GATA6, CLDN18, TFF1, HNF1A, HNF4A) and 4 basal-like markers (S100A2, KRT17, MUC16, PanBS) were selected because they were highly expressed and specific to the corresponding subtype.

The first step of selection on transcriptomic data was made from bulk analyses mixing RNAs from tumor and stromal cells and was also potentially influenced by the tumor cellularity. To address these issues, we used transcriptomic data from 30 murine xenografts of human PDAC (PDX) in which stromal (murine) and tumor (human) expression level could be distinguished(13). We assessed the expression of selected genes in the tumor compartment according to the basal-like and classical molecular subtypes (Supplementary Figure 3). The 9 “purified” markers were indeed differentially expressed according to the molecular subtypes in PDX (p<0.05).

The final step was to validate the selected markers in a large set of 50 human resected PDACs. As tumor subtypes are based on RNA levels, we evaluated the concordance between the IHC expression and the corresponding mRNA level obtained by RNAseq (Supplementary Figure 4). IHC and RNAseq were performed on the same tumor area on the same tissue block. There was a strong and significant correlation (p<0.001) between the IHC expression level and the mRNA transcript level for TFF1 (r=0.87), CLDN18 (r=0.82), S100A2 (r=0.78), KRT17 (r=0.74), PanBS (r=0.62), MUC16 (r=0.59) and GATA6 (r=0.56). The association was non-significant for HNF4A (r=0.19, p=0.2) and HNF1A (r=0.18, p=0.22) and led us to remove these markers. Finally, the validated markers were GATA6, CLDN18, TFF1 for the classical subtype and MUC16, S100A2, KRT17, PanBS for the basal-like subtype.

Overall, this multi-step process allowed us to have a thorough and rational selection of markers to differentiate the classical and basal-like subtypes of a tumor. Our final selection of markers showed a clear anti-correlation between the two groups within this cohort (Figure 1D). However, while very specific, not a single marker has enough sensibility to be used alone to define the two molecular subtypes (Figures 1B-C). This observation reinforces the perspective of a non-dichotomous classification and suggests an intratumoral heterogeneity, which our marker panel might help to explore.

### Four types of tumoral clusters are uncovered using an unsupervised tile clustering

To assess the intratumor heterogeneity by the spatial co-expression of the selected markers, we aligned digitized whole-slide images corresponding to the H&E section and the IHCs (PANCK, GATA6/S100A2, CLDN18/KRT17, TFF1/PanBs, MUC16) (Figure 2A). We used this approach massively on a cohort of 95 patients with resected PDAC. Tumor regions were manually annotated by an expert pathologist (JC) on HES images. Each annotation was then copied on the serial IHC slides and was virtually cut into 200 µm square tiles registered (*i.e.* aligned) on each slide. Positively stained pixels (PANCK, TFF1, CLDN18, GATA6, KRT17, MUC16, PanBs and S100A2) were quantified to obtain a scoring proportion of each marker for each tile. The 106,853 tiles coming from all patients were filtered in a two-step process in order to keep first the non-truncated (85,564) tiles and then the tumor (44,024) tiles containing more than 10% of PanCK staining.

**Figure 2.**
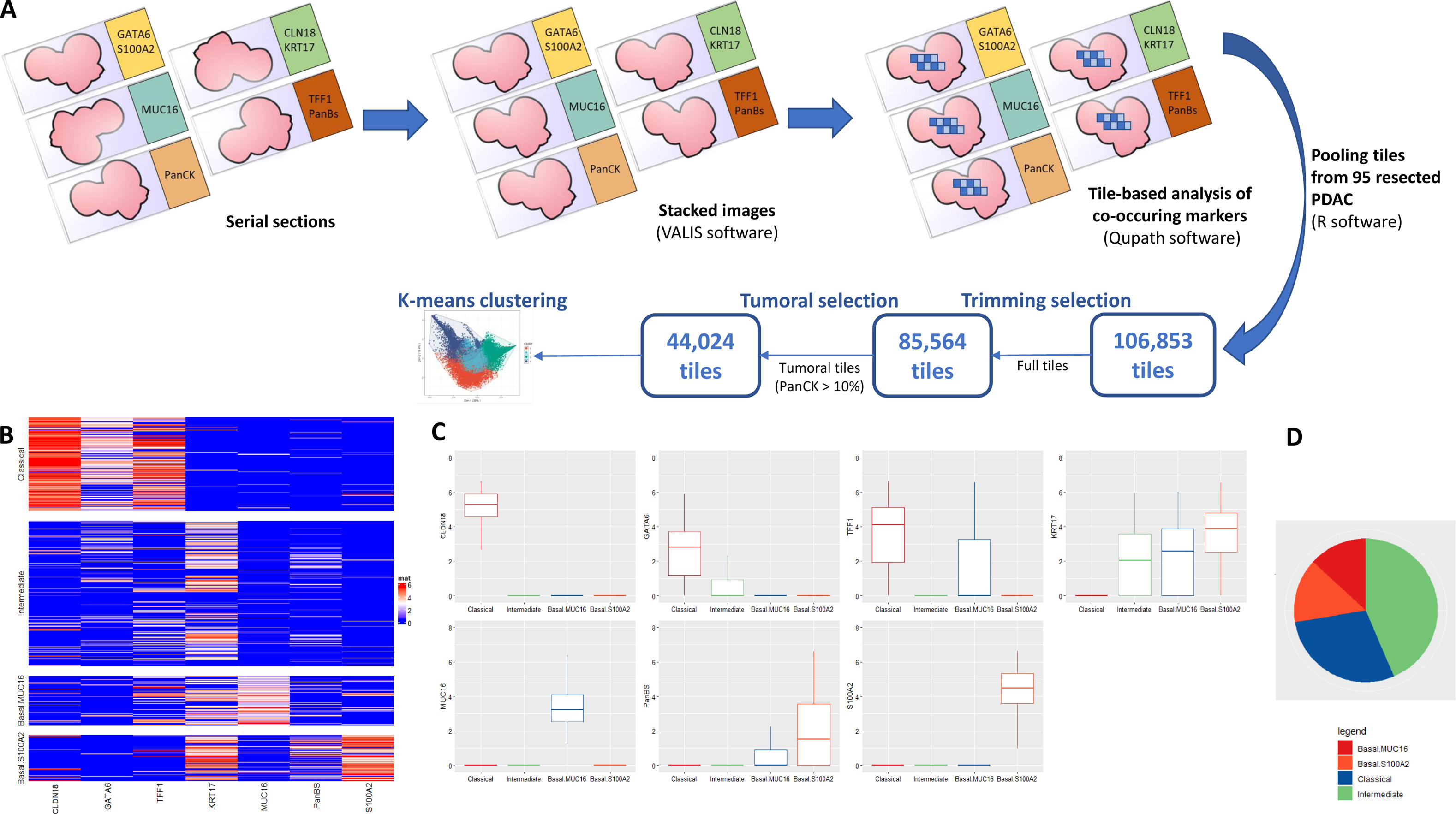
Tile clustering to explore pancreatic intratumoral heteterogeneity. A. Stacked multiplex immunohistochemistry for intratumoral subtype discovery using a tile-based approach. PDAC: pancreatic ductal adenocarcinoma B. Heatmap representing marker expression, as measured by the proportion of positive pixels per per tile (log2 transformed), according to clusters. Rows are tumor tiles (n=44,024). C. Boxplots representing marker expression according to clusters. D. Pie chart representing proportions of clusters in whole slides from 95 resected pancreatic ductal adenocarcinoma.

To validate our marker panel, we evaluated the correlation of each marker with the classical and basal definition according to Moffitt(20) (Supplementary Figure 5). Classical markers significantly correlated with the classical signature and anticorrelated with the basal signature (p<0.05), and *vice versa* for basal markers except for MUC16 that did not significantly anticorrelate with the classical signature (p=0.30) but strongly correlated with the basal signature (p<0.0001). This observation encouraged us to pursue further with our marker panel.

K-means clustering of tiles using the proportion of positive pixels of the 7 subtype markers revealed four types of tiles (Supplementary Figure 6). Clusters according to marker expression are shown in Figures 2B-C. We found two basal clusters including one with a predominant expression of MUC16, named Basal. MUC16, and one with a predominant expression of S100A2, named Basal.S100A2. A third cluster was typically Classical with a mixed expression of CLDN18, GATA6 and TFF1 and the fourth cluster was negative for all markers except for KRT17 and to a lesser extent for GATA6 and was named Intermediate. By pooling all tiles, basal, classical and intermediate clusters represented 27.3%, 27.4% and 45.3% respectively (Figure 2D). Examples of representative areas of each cluster are depicted in Supplementary Figure 7.

To validate these findings, we correlated each cluster expression with the previously defined molecular signatures (Figure 3A). Basal.S100A2 and Basal.MUC16 expression correlated with basal-like signatures (Puleo, Moffit, Bailey, Purist Score) (p<0.001). In contrast, the Classical cluster correlated with classical signatures (p<0.001) and the Intermediate cluster did not correlate with neither basal nor classical signatures (p>0.05). The expression of clusters in the 95 PDACs according to the Purist score is represented in Figure 3B and the proportion of each cluster by their major component is shown in Figure 3C. The Intermediate cluster was almost always present (96%) and was predominant in 41% of PDACs. The mix of classical, basal and intermediate clusters was present in 62.3% of cases highlighting the important and constant intratumoral heterogeneity of PDAC. To explore the spatial conservation of this heterogeneity, we analyzed spatially distinct areas from 2 different blocks from the same PDAC in 23 patients (Figure 3D). Interestingly, the predominant cluster remained the same for 16 PDAC (70%) highlighting the conserved heterogeneity in space.

**Figure 3.**
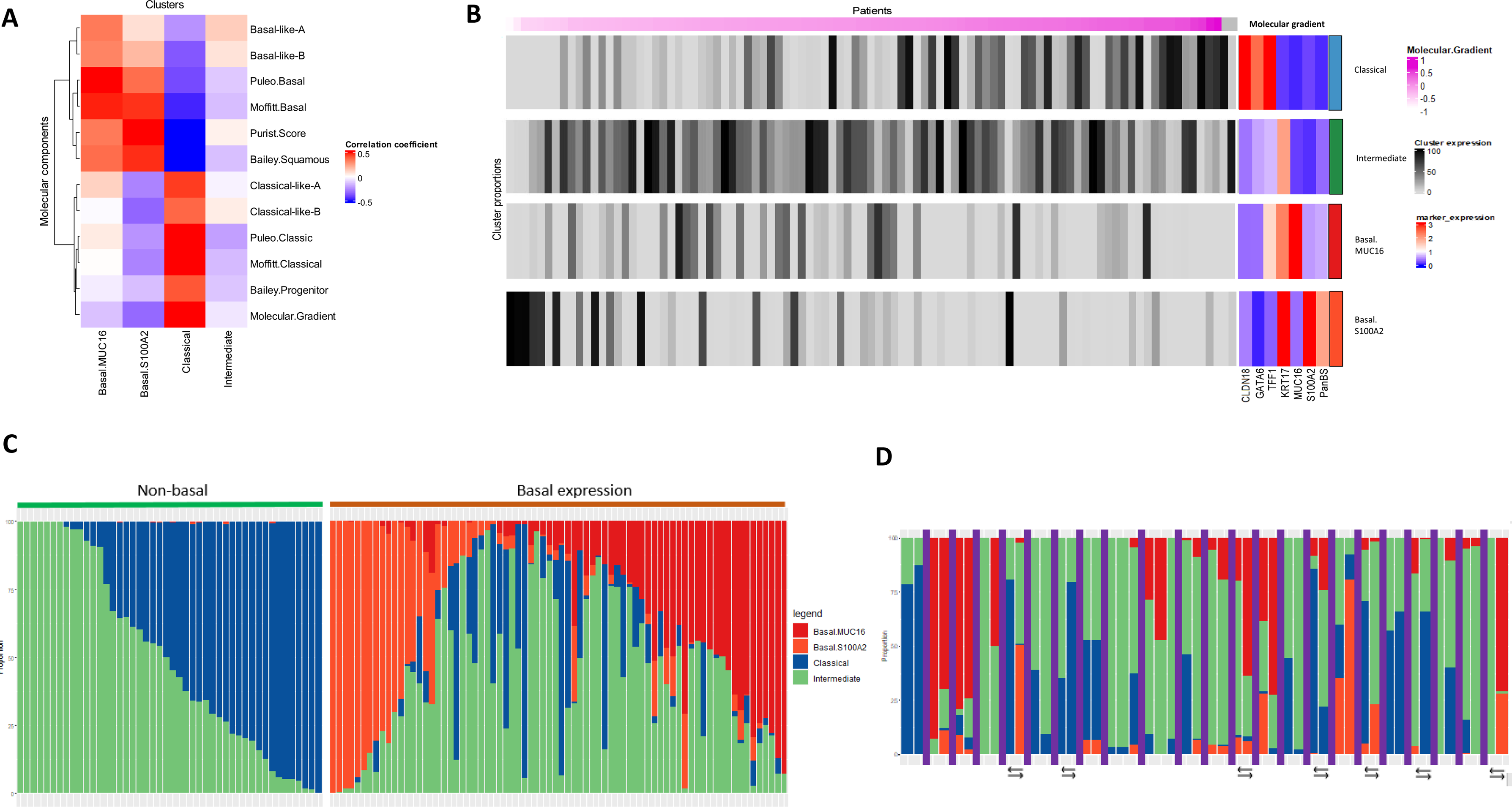
Representation of tile clusters and association with the molecular subtypes. A. Heatmap representing correlation of tile-cluster proportions with transcriptomic-based subtypes. B. Heatmap representing the proportion of each cluster according to the molecular gradient. Mean marker expression according to each cluster is represented on right side of the heatmap. C. Bar plots representing tile-cluster proportions in each tumor. Tumors are split between basal expression (> 1% of tiles) and non-basal expression (≤ 1% of tiles). D. Bar plots representing per-slide tile-cluster proportions for each tumor with multiple slides (n=23). Slides from different tumors are separated by a purple vertical bar. Arrows represent shift in predominant cluster in the same tumor.

### The presence of a basal contingent sharply reduces survival, regardless of the remaining PDAC composition

To assess the importance of assessing intratumoral hetereogeneity, we then evaluated whether the cluster expression among the 95 patients had a prognostic impact. Neither the presence (>1%) nor the predominance (>50%) of the Intermediate cluster did impact OS (p>0.50). Only the predominance of the Classical cluster was associated with a better OS (p=0.045) but not significantly in the multivariate analysis (HR=0.59, p=0.09). In contrast, patients with PDAC containing even a minor basal (>1%) component had a strong decreased OS in univariate (median OS 30.5 months vs 62.8 months, p=0.002) and multivariate analyses (HR=2.36, p=0.01) (Figure 4A-B). This prognostic association was also observed in an independent cohort of 148 patients with PDAC on TMA (H-score > 0 regarding S100A2 or MUC16) in univariate (median OS 33.1 months vs 50.6 months, p=0.04) and multivariate analyses (HR=1.61, trend with p=0.07) (Figure 4C). Of note, TMA captures only one part of the tumor, and this could explain the trend rather than the significance by missing basal-like cells in some cases.

**Figure 4.**
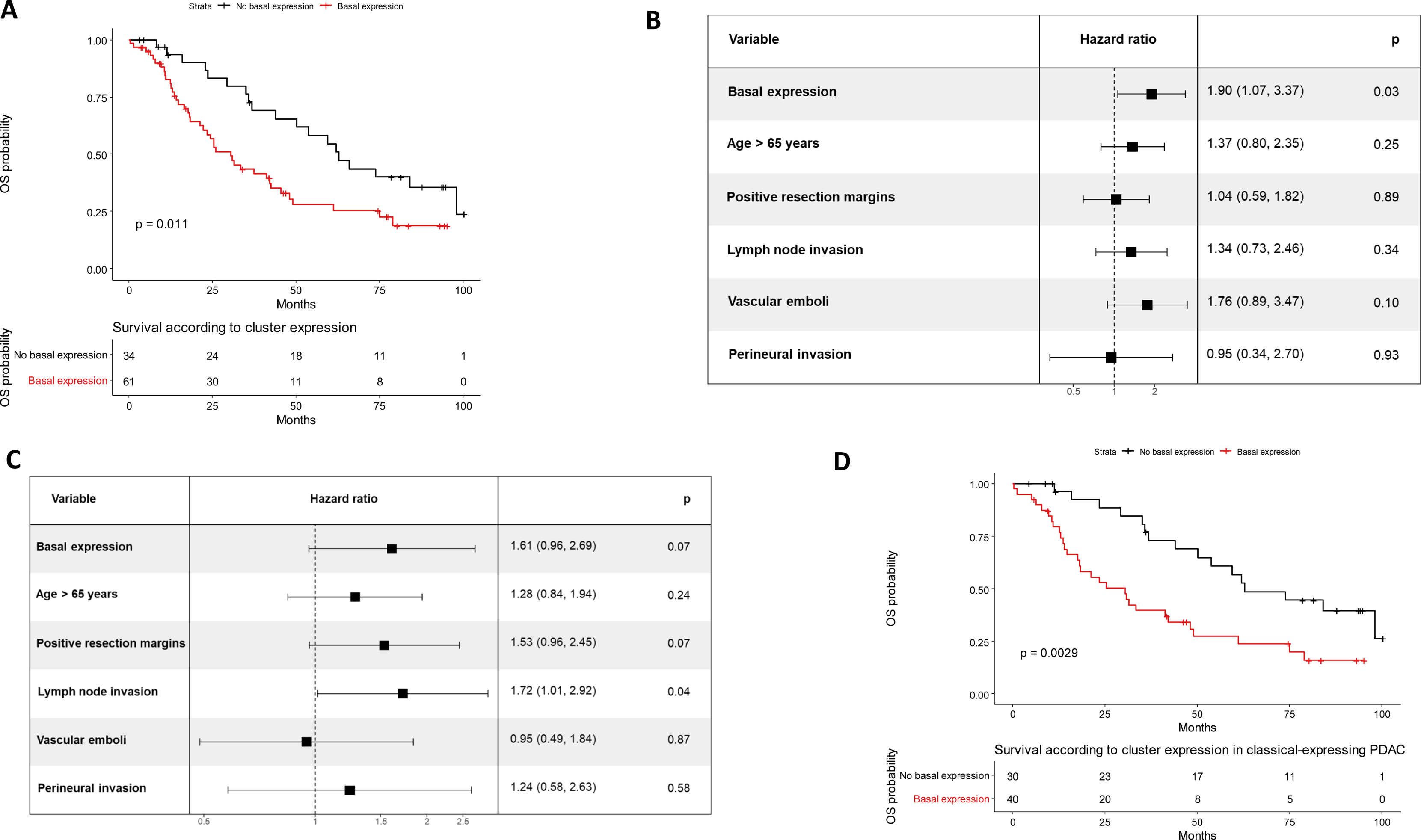
Survival of patients with PDAC according to the cluster expression. A. Overall survival curves according to the detection of basal-expressing tiles (>1% of tiles) in patients with PDAC (n=95 patients). OS: overall survival. B. Multivariate analysis for the detection of basal-expressing tiles (>1% of tiles) and clinicopathological factors regarding overall survival (n=95 patients). C. Multivariate analysis for the basal expression (H-score MUC16 or S100A2 > 0) and clinicopathological factors regarding overall survival (n=148 patients). D. Overall survival curves according to the basal expression (>1% of tiles) in patients with classical-expressing (>1% of tiles) PDAC (n=70 patients). OS: overall survival.

In addition, when restricting the analysis to patients with PDAC expressing the Classical cluster (>1%), the observed difference in OS was even greater according to the co-occurence of a basal component (p<0.001) (Figure 4D). Indeed, the group of patients with classical but no basal expression had a much longer OS in multivariate analysis (HR=0.34, p=0.003).

By exploring each basal cluster independently, we found that the Basal.MUC16 cluster expression significantly and negatively impacted OS while a trend toward a decreased OS was observed for Basal.S100A2 cluster expression in multivariate analyses (HR=1.93, p=0.02; HR=1.63, p=0.08, respectively) (Supplementary Figure 8A-B). Among patients with basal expression, OS was not impacted by the predominance of one of the basal clusters (p=0.64) (Supplementary Figure 8C). Similarly, OS among patients without basal expression was not impacted by the predominance of the classical or intermediate clusters (p=0.91) (Supplementary Figure 8D). Besides, none of the traditional clinico-pathological parameters among T and N status, tumor differentiation, resection margins, vascular and perineural invasion were significantly associated with a cluster expression (p>0.05).

Overall, the presence of the basal phenotype was the most important prognostic factor, even in PDAC expressing the classical cluster.

### The evolutionary process of pancreatic carcinogenesis may start from a classical toward a basal phenotype by going through an intermediate phenotype

Given that the previous steps have shown the relevance of our marker panel, we wanted to explore which clusters are present at the first stages of pancreatic carcinogenesis. To do so, we studied the markers expression (h-score) in preneoplastic lesions among 16 PDACs (4 classical-predominant, 4 basal-predominant and 8 intermediate-predominant) including 5 early acinar-to-ductal metaplasia (ADM), 6 late ADM, 7 low-grade pancreatic intraepithelial neoplasias (PanINs) and 7 high-grade PanINs. Fifteen normal areas were also analyzed (normal duct and centroacinar cells). Normal duct and centroacinar cells, and early ADM strongly and exclusively expressed GATA6 (Figure 5A-B). Surprisingly, a moderate expression of KRT17 appeared in late ADMs and high-grade PanINs along with a decrease in expression in classical markers in their early stage (early ADMs and low-grade PanINs), independently of the basal expression in the adjacent PDAC. Other basal markers were weakly expressed or absent from normal and preneoplastic lesions. When expressed in PanINs, basal markers (KRT17, S100A2 and PanBS) were located at the basal cell pole whereas classical markers (CLDN18, GATA6, TFF1) were at the apical pole (Supplementary Figure 9). Altogether, these findings indicate the presence of a classical phenotype and KRT17, as an early basal marker, in preneoplastic lesions. Specific markers of basal clusters (i.e. MUC16 and S100A2) appear later during the PDAC evolution, making them late basal markers.

**Figure 5.**
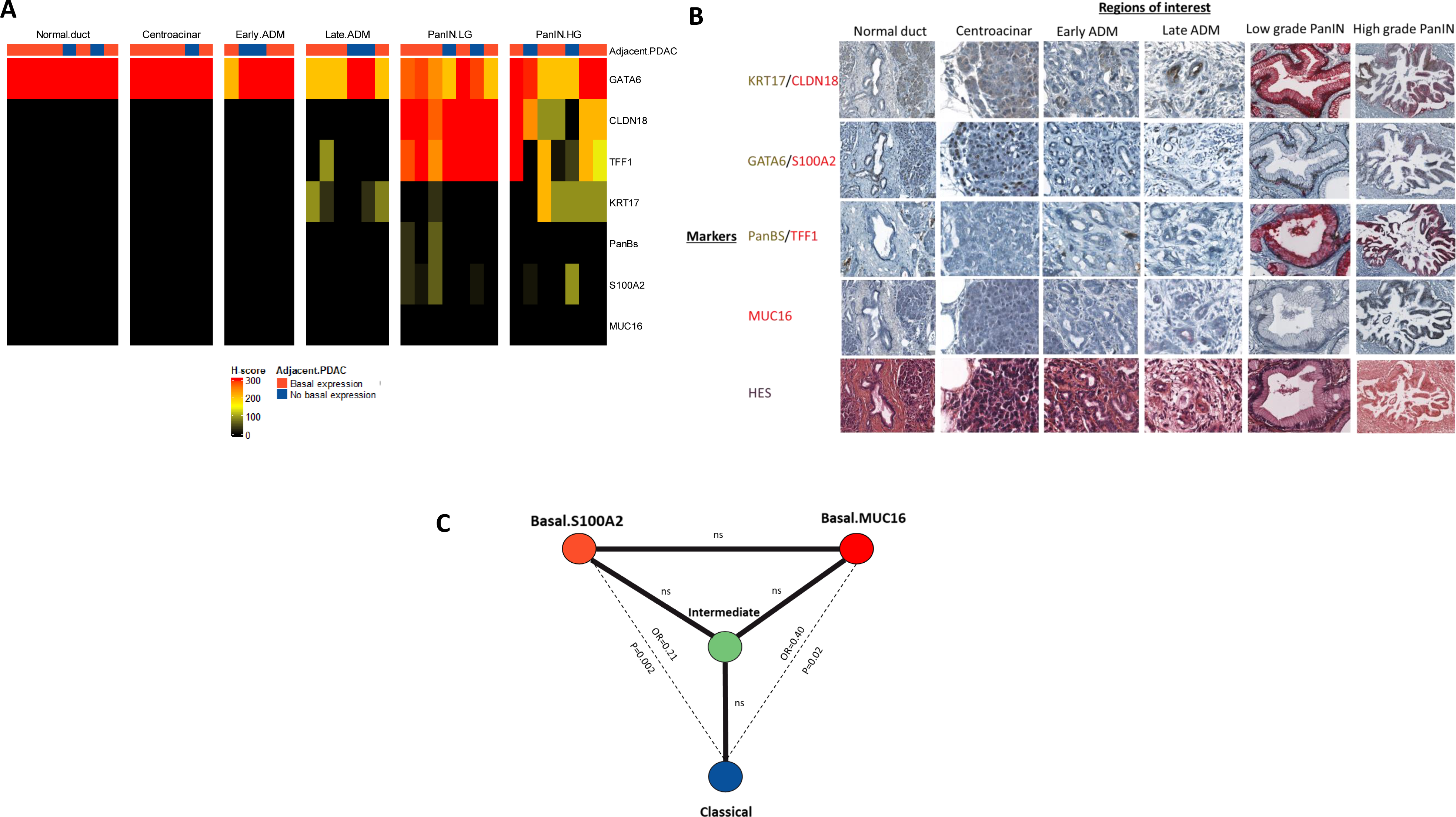
Marker expression in preneoplastic lesions and co-occurrence between clusters. A. Heatmap representing marker expression in normal cells and preneoplastic lesions. ADM: acinar-to-ductal metaplasia; HG: high-grade; LG: low-grade; PanIN: pancreatic intraepithelial neoplasia. H-score: 0 (black) to 300 (red). B. Marker expression according to regions of interest in representative areas. ADM: acinar-to-ductal metaplasia; HG: high-grade; LG: low-grade; PanIN: pancreatic intraepithelial neoplasia. C. Clusters co-occurrence in 95 resected PDAC (Fisher test).

At this stage, our hypothesis was that the Intermediate cluster with the predominant expression of KRT17 represents the missing link in a continuum between the classical and the basal tumor cell phenotypes. By analyzing the co-occurrence of phenotypes in each tumor, we observed that the intermediate cluster co-occurred frequently with the other phenotypes, contrary to basal-like and Classical clusters (Figure 5C). This suggested that the intermediate cells may be a transitional state between the classical and the basal cluster. Furthermore, 6 out of the 7 shifts (*i.e.* change in the predominant phenotype) observed in PDACs with more than one slide (n=23) occurred between the Intermediate and another cluster, and one between the Classical and basal clusters (Figure 3D), reinforcing our starting hypothesis.

As co-occurrence between both basal clusters was not less frequent than expected by chance (Figure 5C), we suspected 2 differential steps of evolution. To understand the biological substratum between the 2 basal clusters, we explored their differences. Firstly, we observed that the 6 adenosquamous cases were all Basal.S100A2 predominant, while none expressed more than 2% of Basal.MUC16. We then compared gene expressions of PDACs with a major expression of Basal.MUC16 and less than 5% of Basal.S100A2 (n=5) versus PDACs with a major expression of Basal.S100A2 and less than a 5% of Basal.MUC16 (n=6, including 4 out of 6 adenosquamous cases). Firstly, genes involved in the classical program were decreased in the Basal.S100A2 cluster compared to the Basal.MUC16 cluster (Supplementary Figure 10A), consistently with the previous correlations (Figure 3A) and suggesting a more distant state of differentiation. Secondly, the Basal.MUC16 cluster showed a significant increase (adjusted p<0.05) in genes associated with bacterial infection (*PDZD3, SFTPA2, PIK3C2B*), cytokine signaling (*TRIM31, SERPINB2*) and apoptosis (BCL2L15, VIL1). The Basal.S100A2 cluster significantly upregulated genes associated with nervous system development (IRX3, KRT6B, PKP1, ABC4, DSC3), ECM modulation (MMP13, COL7A1, PAK6), and cell proliferation (FAT2, S1PR5, GPC1, PTTG1) (Supplementary Figure 10B). Gene set enrichment analysis validated the significant upregulation of proliferative pathways in the Basal.S100A2 cluster and inflammatory pathways in Basal.MUC16 cluster (Supplementary Figure 10A).

## Discussion

Herein, we developed through this work a panel of antibodies that could be easily used by researchers and pathologists. The purpose of this panel was to classify patients according to the two main subtypes of PDAC, roughly basal-like or classical. To achieve this, we selected markers through a stringent and multiple process. These markers had to be associated with either classical and basal components, have functional antibodies, be specific to tumor cells and be sufficiently expressed in human tissues. This highlighted the limit of RNA-based selection as some genes such as *KRT6A* were linked to the epithelial component, regardless of the subtype, but appeared as strongly differential due to the tumor cellularity of basal-like tumors being much higher than classical tumors. Current scientific knowledge of PDAC suggests that this cancer presents a significant inter-tumoral and intra-tumoral heterogeneity. Our panel of 7 markers (GATA6, CLDN18, TFF1, MUC16, S100A2, KRT17, PanBS) uncovered the limits of the binary basal-like/classical classification by highlighting the major Intermediate cluster co-expressing classical and basal-like markers, which does not fit into this classification. Besides, by registering serial whole IHC slides and performing a tile-based analysis, we demonstrated a high degree of intratumoral heterogeneity, with a frequent co-occurrence of classical, basal and intermediate clusters, both between tumors and within the same tumor. This tremendous intratumoral heterogeneity and likely subsequent plasticity could explain the treatment resistance and the very poor prognosis of PDAC.

Our study shows the difficulty of the interpretation of PDAC heterogeneity using bulk transcriptomic approaches, and thus limiting their routine application. In addition, analysis takes longer than IHC interpretation, requires the availability of sequencing facilities and RNA-based approaches are limited by the quantity and quality of samples (e.g., formalin fixation or low cellularity, as in biopsies). Finally, RNAseq is expensive when performed on a low number of samples and requires important normalization from batch to batch. Several studies have already identified biomarkers for routine tumor subtyping but were limited to few markers (16, 25, 26). The strength of our study was to thoroughly select routine pathology-grade surrogate markers, and to analyze quantifications using a tile-based approach on stacked slides. This process allows to perform multiplex-like whole slide IHC on large cohorts, on routine pathology automates and for a fraction of the cost of state-of-the-art techniques such as CODEX or LUNAPHORE. It should be noted that our approach performed on serial slides allows the spatial phenotyping of a group of cells but not the exact complete phenotyping of single cells. Through this different method, we highlighted the potential of few IHC markers to optimally define intratumoral PDAC heterogeneity. Williams et al. recently proposed a multiplex immunofluorescence panel including 6 of our markers(11) (CLDN18.2, TFF1, GATA6, KRT17, KRT5, and S100A2). Given the strong prognostic value of the basal expression and the distinct biology of the Basal.MUC16 cluster, adding MUC16 would be optimal to entirely capture the basal expression. Facilitating the assessment of tumor heterogeneity could also be useful in the case of sequential tumor assessment under experimental treatment modulating PDAC phenotype (e.g. epigenetic strategies)(27).

Overall, our thorough selection of basal and classical markers classified 54.7 % of tumoral tiles as basal or classical but the remaining tiles had a non-classical and a non-basal phenotype. The intermediate phenotype was characterized by the predominant expression of KRT17, non-specific of the basal subtype. This confirmed its intermediate nature, as reported in the single cell study by Williams et al.(11), in which KRT17 was activated in cells that co-expressed basal and classical markers. Our work also highlighted that not all markers are expressed similarly throughout the classical-intermediate-basal transition. Claudin 18 was the most restrictive marker, exceptionally co-expressed with KRT17 in the same tile while TTF1 (rarely) and more frequently GATA6 could be co-expressed with KRT17 and sometimes S100A2 or MUC16. KRT17 could be expressed alone or with the late basal markers S100A2 and MUC16 that were exclusive of classical markers. Consistently with Williams et al., the intermediate state was almost always present in PDAC, showing its potential importance in the transition between classical and basal-like subtypes. By analyzing preneoplastic lesions, we showed evidence of the evolution process of pancreatic carcinogenesis going from a classical toward a basal phenotype through an intermediate phenotype. This is consistent with the observation that the final stage of PDAC development (i.e. metastases) is dominated by basal cells(11). Sequential samples would help support this hypothesis and the exact mechanisms of this transition (e.g. epigenetic remodeling) remains to be defined.

Importantly, our study brings new insight into the presence and the possible co-occurrence of 2 different basal phenotypes. Basal.MUC16 and Basal.S100A2 clusters had specific expressions of MUC16 and S100A2, respectively. Consistently, both markers have been shown to be associated with poor prognosis in previous studies(28, 29). The Basal.MUC16 cluster was more associated with cytokine signaling whereas the Basal.S100A2 cluster was more proliferative and particularly far from the classic phenotype. It is well described that basal-like tumors relapse more often after surgery(30) and are less responsive to chemotherapy(16, 31). Although the prognostic difference between both basal phenotypes was not apparent in our study, the information that a minor expression of one of two was enough to drastically reduce OS is important and coherent with our previous work(2). This highlights the importance of a complete tumor analysis, as some classical PDACs may harbour aggressive basal contingents. One limitation of our study is that it is based on a two-dimensional approach that does not allow exhaustive exploration of the tumor. The development of 3D approaches may help overcome this issue in the future. Nevertheless, our IHC panel redefined and easily assessed the high degree of intra- and inter-tumoral heterogeneity of PDAC.

## References

1. Integrated Genomic Characterization of Pancreatic Ductal Adenocarcinoma. Cancer Cell. 2017;32(2):185–203.e13.

2. Saillard C, Delecourt F, Schmauch B, Moindrot O, Svrcek M, Bardier-Dupas A, et al. Pacpaint: a histology-based deep learning model uncovers the extensive intratumor molecular heterogeneity of pancreatic adenocarcinoma. Nature communications. 2023;14(1):3459.

3. Chan-Seng-Yue M, Kim JC, Wilson GW, Ng K, Figueroa EF, O’Kane GM, et al. Transcription phenotypes of pancreatic cancer are driven by genomic events during tumor evolution. Nat Genet. 2020;52(2):231–40.

4. Topham JT, Karasinska JM, Lee MKC, Csizmok V, Williamson LM, Jang GH, et al. Subtype-Discordant Pancreatic Ductal Adenocarcinoma Tumors Show Intermediate Clinical and Molecular Characteristics. Clin Cancer Res. 2021;27(1):150–7.

5. Yachida S, Jones S, Bozic I, Antal T, Leary R, Fu B, et al. Distant metastasis occurs late during the genetic evolution of pancreatic cancer. Nature. 2010;467(7319):1114-7.

6. Cros J, Raffenne J, Couvelard A, Poté N. Tumor Heterogeneity in Pancreatic Adenocarcinoma. Pathobiology. 2018;85(1-2):64–71.

7. S NK, Wilson GW, Grant RC, Seto M, O’Kane G, Vajpeyi R, et al. Morphological classification of pancreatic ductal adenocarcinoma that predicts molecular subtypes and correlates with clinical outcome. Gut. 2020;69(2):317–28.

8. Hayashi A, Fan J, Chen R, Ho YJ, Makohon-Moore AP, Lecomte N, et al. A unifying paradigm for transcriptional heterogeneity and squamous features in pancreatic ductal adenocarcinoma. Nat Cancer. 2020;1(1):59–74.

9. Nicolle R, Blum Y, Duconseil P, Vanbrugghe C, Brandone N, Poizat F, et al. Establishment of a pancreatic adenocarcinoma molecular gradient (PAMG) that predicts the clinical outcome of pancreatic cancer. EBioMedicine. 2020;57:102858.

10. Juiz N, Elkaoutari A, Bigonnet M, Gayet O, Roques J, Nicolle R, et al. Basal-like and classical cells coexist in pancreatic cancer revealed by single-cell analysis on biopsy-derived pancreatic cancer organoids from the classical subtype. FASEB J. 2020;34(9):12214–28.

11. Williams HL, Dias Costa A, Zhang J, Raghavan S, Winter PS, Kapner KS, et al. Spatially Resolved Single-Cell Assessment of Pancreatic Cancer Expression Subtypes Reveals Co-expressor Phenotypes and Extensive Intratumoral Heterogeneity. Cancer Res. 2023;83(3):441–55.

12. Puleo F, Nicolle R, Blum Y, Cros J, Marisa L, Demetter P, et al. Stratification of Pancreatic Ductal Adenocarcinomas Based on Tumor and Microenvironment Features. Gastroenterology. 2018;155(6):1999–2013.e3.

13. Nicolle R, Blum Y, Marisa L, Loncle C, Gayet O, Moutardier V, et al. Pancreatic Adenocarcinoma Therapeutic Targets Revealed by Tumor-Stroma Cross-Talk Analyses in Patient-Derived Xenografts. Cell Rep. 2017;21(9):2458–70.

14. Gatenbee CD, Baker A-M, Prabhakaran S, Swinyard O, Slebos RJC, Mandal G, et al. Virtual alignment of pathology image series for multi-gigapixel whole slide images. Nature communications. 2023;14(1):4502.

15. Bankhead P, Loughrey MB, Fernández JA, Dombrowski Y, McArt DG, Dunne PD, et al. QuPath: Open source software for digital pathology image analysis. Sci Rep. 2017;7(1):16878.

16. O’Kane GM, Grünwald BT, Jang GH, Masoomian M, Picardo S, Grant RC, et al. GATA6 Expression Distinguishes Classical and Basal-like Subtypes in Advanced Pancreatic Cancer. Clin Cancer Res. 2020;26(18):4901–10.

17. Collisson EA, Sadanandam A, Olson P, Gibb WJ, Truitt M, Gu S, et al. Subtypes of pancreatic ductal adenocarcinoma and their differing responses to therapy. Nat Med. 2011;17(4):500–3.

18. Bailey P, Chang DK, Nones K, Johns AL, Patch AM, Gingras MC, et al. Genomic analyses identify molecular subtypes of pancreatic cancer. Nature. 2016;531(7592):47–52.

19. Kalisz M, Bernardo E, Beucher A, Maestro MA, Del Pozo N, Millán I, et al. HNF1A recruits KDM6A to activate differentiated acinar cell programs that suppress pancreatic cancer. EMBO J. 2020;39(9):e102808.

20. Moffitt RA, Marayati R, Flate EL, Volmar KE, Loeza SGH, Hoadley KA, et al. Virtual microdissection identifies distinct tumor- and stroma-specific subtypes of pancreatic ductal adenocarcinoma. Nat Genet. 2015;47(10):1168–78.

21. Rashid NU, Peng XL, Jin C, Moffitt RA, Volmar KE, Belt BA, et al. Purity Independent Subtyping of Tumors (PurIST), A Clinically Robust, Single-sample Classifier for Tumor Subtyping in Pancreatic Cancer. Clin Cancer Res. 2020;26(1):82–92.

22. Somerville TDD, Xu Y, Miyabayashi K, Tiriac H, Cleary CR, Maia-Silva D, et al. TP63-Mediated Enhancer Reprogramming Drives the Squamous Subtype of Pancreatic Ductal Adenocarcinoma. Cell Rep. 2018;25(7):1741–55.e7.

23. Roa-Peña L, Leiton CV, Babu S, Pan CH, Vanner EA, Akalin A, et al. Keratin 17 identifies the most lethal molecular subtype of pancreatic cancer. Sci Rep. 2019;9(1):11239.

24. Thomas D, Sagar S, Liu X, Lee HR, Grunkemeyer JA, Grandgenett PM, et al. Isoforms of MUC16 activate oncogenic signaling through EGF receptors to enhance the progression of pancreatic cancer. Mol Ther. 2021;29(4):1557–71.

25. Muckenhuber A, Berger AK, Schlitter AM, Steiger K, Konukiewitz B, Trumpp A, et al. Pancreatic Ductal Adenocarcinoma Subtyping Using the Biomarkers Hepatocyte Nuclear Factor-1A and Cytokeratin-81 Correlates with Outcome and Treatment Response. Clin Cancer Res. 2018;24(2):351–9.

26. Wang Y, Park JYP, Pacis A, Denroche RE, Jang GH, Zhang A, et al. A Preclinical Trial and Molecularly Annotated Patient Cohort Identify Predictive Biomarkers in Homologous Recombination-deficient Pancreatic Cancer. Clin Cancer Res. 2020;26(20):5462–76.

27. Versemann L, Hessmann E, Ulisse M. Epigenetic Therapeutic Strategies to Target Molecular and Cellular Heterogeneity in Pancreatic Cancer. Visc Med. 2022;38(1):11–9.

28. Muniyan S, Haridas D, Chugh S, Rachagani S, Lakshmanan I, Gupta S, et al. MUC16 contributes to the metastasis of pancreatic ductal adenocarcinoma through focal adhesion mediated signaling mechanism. Genes Cancer. 2016;7(3-4):110–24.

29. Chen Y, Wang C, Song J, Xu R, Ruze R, Zhao Y. S100A2 Is a Prognostic Biomarker Involved in Immune Infiltration and Predict Immunotherapy Response in Pancreatic Cancer. Front Immunol. 2021;12:758004.

30. Hilmi M, Cros J, Puleo F, Augustin J, Emile JF, Svrcek M, et al. Tumour and stroma RNA signatures predict more accurately distant recurrence than clinicopathological factors in resected pancreatic adenocarcinoma. Eur J Cancer. 2021;148:171–80.

31. Aung KL, Fischer SE, Denroche RE, Jang GH, Dodd A, Creighton S, et al. Genomics-Driven Precision Medicine for Advanced Pancreatic Cancer: Early Results from the COMPASS Trial. Clin Cancer Res. 2018;24(6):1344–54.

