## Supplementary Data for "Redefining phenotypic intratumor heterogeneity of pancreatic ductal adenocarcinoma: a bottom-up approach"

**Supplementary Table 1: Characteristics of tested antibodies**

| Antibody | Dilution | Clone | Origin | Distributor | Marking location |
|---|---|---|---|---|---|
| c-MET | 1/50 ov | D1C2 | rabbit | Cell Signaling | Cytoplasmic |
| *KRT6A* - CK6A | 1/200 | poly | rabbit | Thermo scientific | Cytoplasmic |
| S100A2 | 1/1000 | EPR5392 | rabbit | abcam | Cytoplasmic, nuclear |
| PanBasal | 1/100 | p63/CK5/CK14 | mouse | Zytomed | Cytoplasmic, membranous, nuclear |
| SNAI | 1/100 | poly | | Santa Cruz | Nuclear |
| *KRT17* - CK17 | 1/50 ov | E3 | mouse | Thermo scientific | Cytoplasmic |
| EGFR | 1/50 ov | EGFR 113 | mouse | novocastra | Membranous |
| *SCL29A1*- ENT | 1/200 | poly | | Spring Bioscience | |
| *SCL16A3* - MCT4 | 1/400 ov | poly | rabbit | Sigma | Membranous |
| MUC16 | 1/250 | X325 | | abcam | Cytoplasmic |
| AGR2 | 1/1000 | D9V2F | rabbit | Cell signaling | Nuclear |
| CYP24A1 | 1/50 | poly | rabbit | Biotechne | |
| HMGA2 | 1/25 ov | D1A7 | rabbit | Cell Signaling | Nuclear |
| GATA6 | 1/200 ov | D61E4 | rabbit | Cell Signaling | Nuclear |
| TFF1 | 1/100 | D2Y1J | rabbit | Cell Signaling | Cytoplasmic |
| CLDN18 | 1/50 | poly | rabbit | Sigma | Cytoplasmic, membranous |
| HNF4A | 1/100 | poly | | sigma | Nuclear |
| REG4 | 1/100 | poly | rabbit | Thermo scientific | |
| HNF1A | 1/50 | poly | rabbit | abcam | Nuclear |
| *KRT20* - CK20 | 1/200 | K020.8 | mouse | DAKO | Membranous |
| CDX2 | 1/750 | EPR2764Y | rabbit | abcam | Nuclear |

**Supplementary Figure 1.** Techniques and aims regarding the 3 cohorts of patients with resected pancreatic ductal adenocarcinoma.

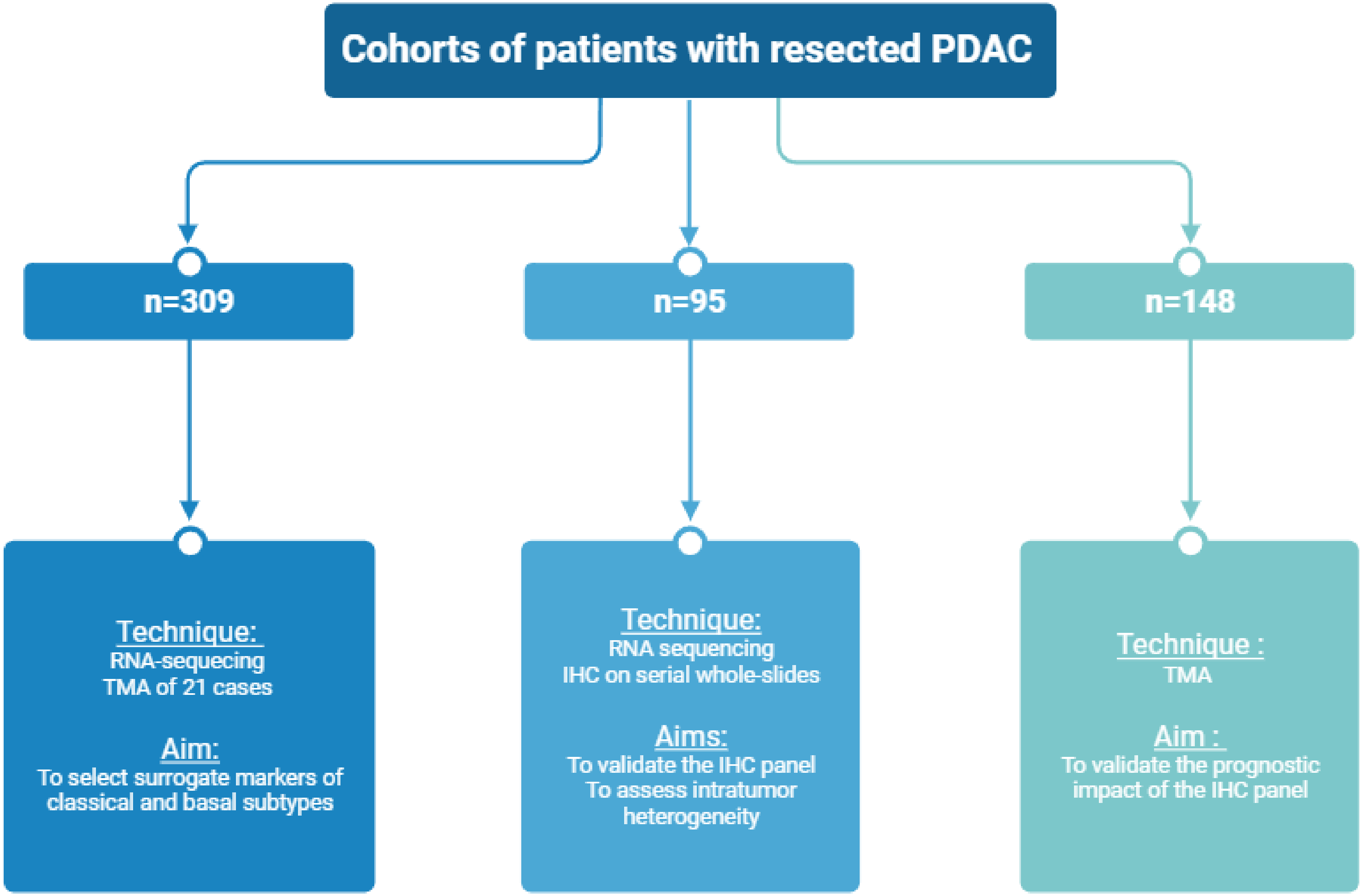

**Supplementary Figure 2.** Example of labelling of the different antibodies tested, with a negative case (H-score=0) on the left panel, an intermediate positive case (H-score=150) in the middle and a strong positive case (H-score=300) on the right.

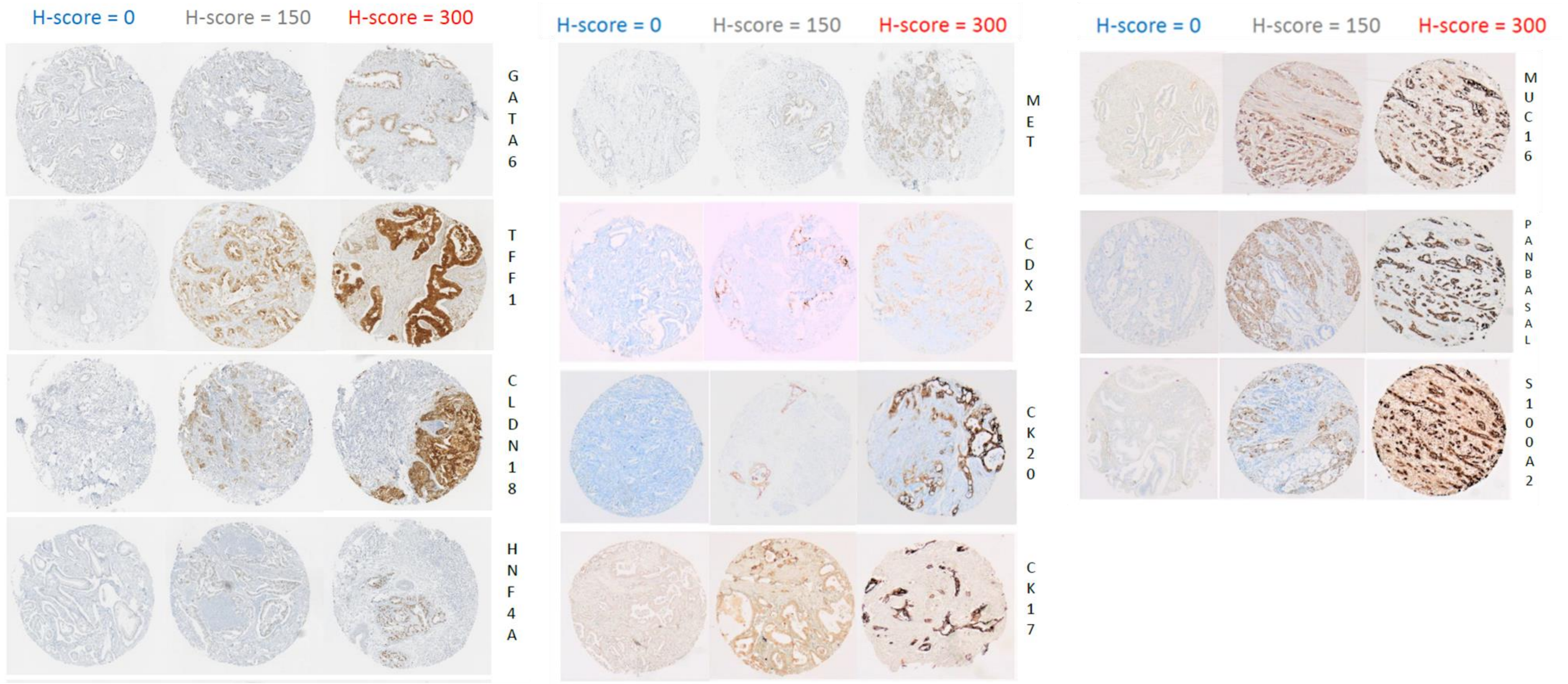

**Supplementary Figure 3.** Expression level of selected genes according to basal-like and classical molecular subtypes in 23 patient-derived xenografts.

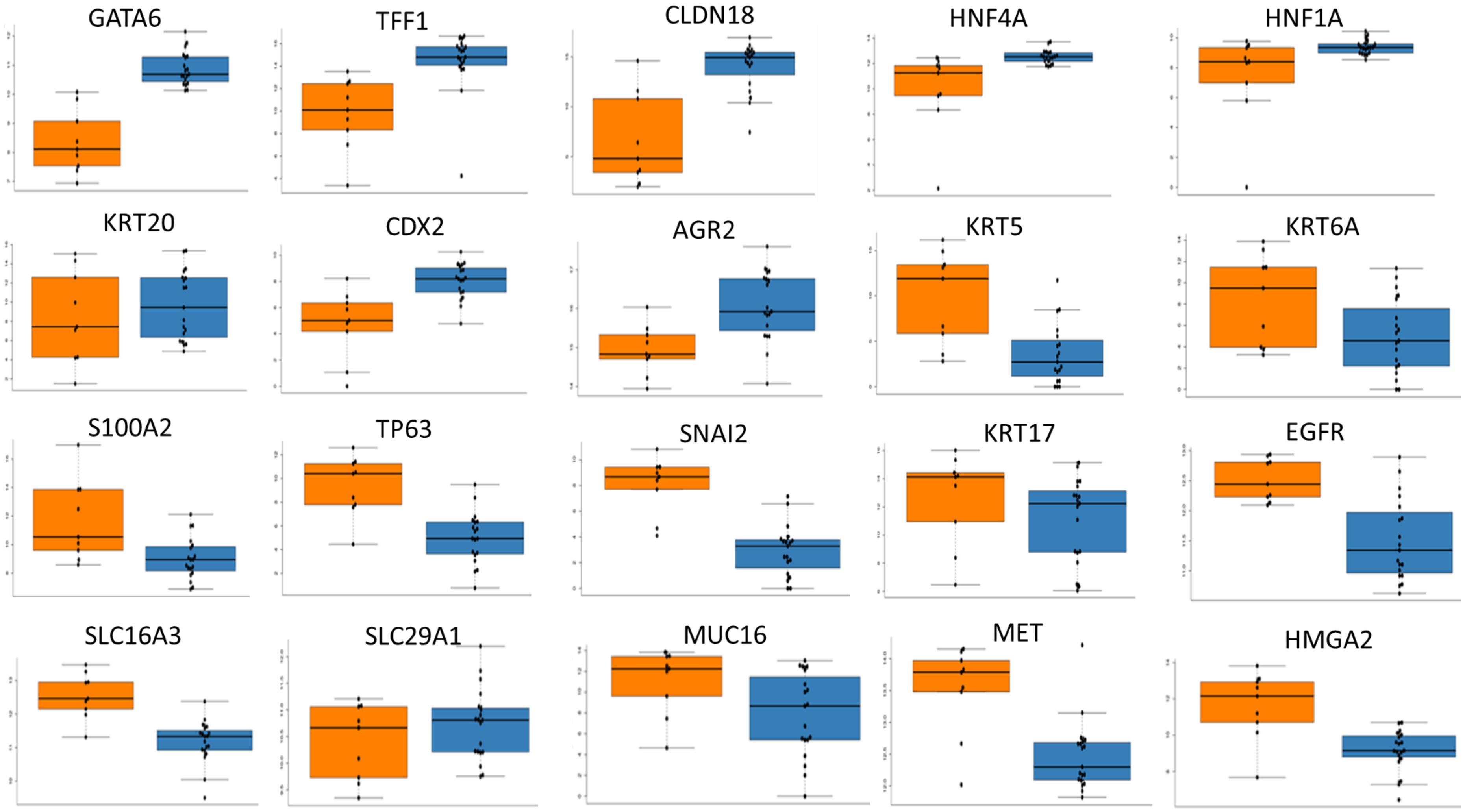

**Supplementary Figure 4.** Correlation between marker expression levels in IHC (H-score) and RNAseq in 50 pancreatic ductal adenocarcinomas.

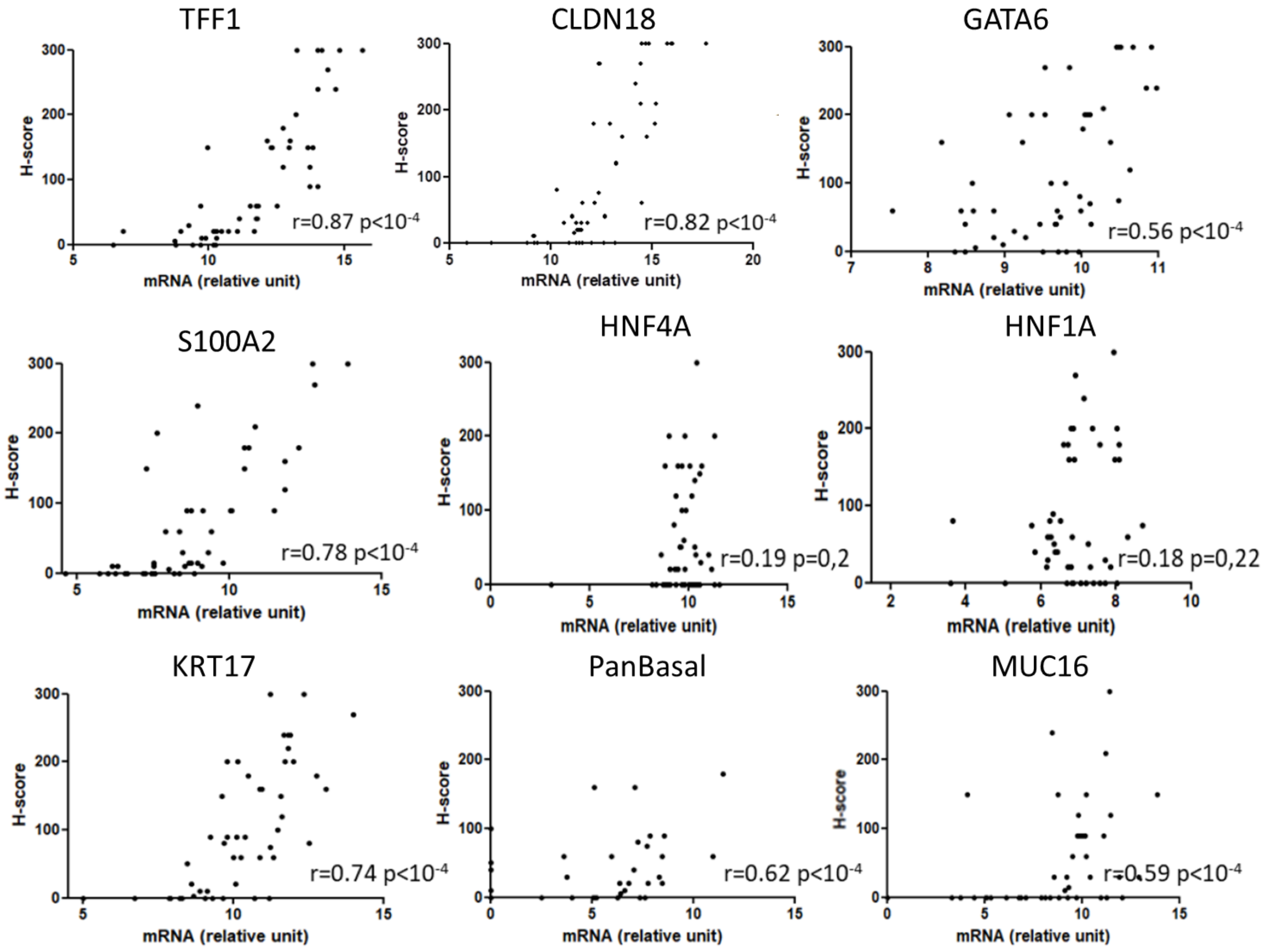

**Supplementary Figure 5.** Correlation of each marker with the classical and basal definition according to Moffitt in 95 pancreatic ductal adenocarcinomas.

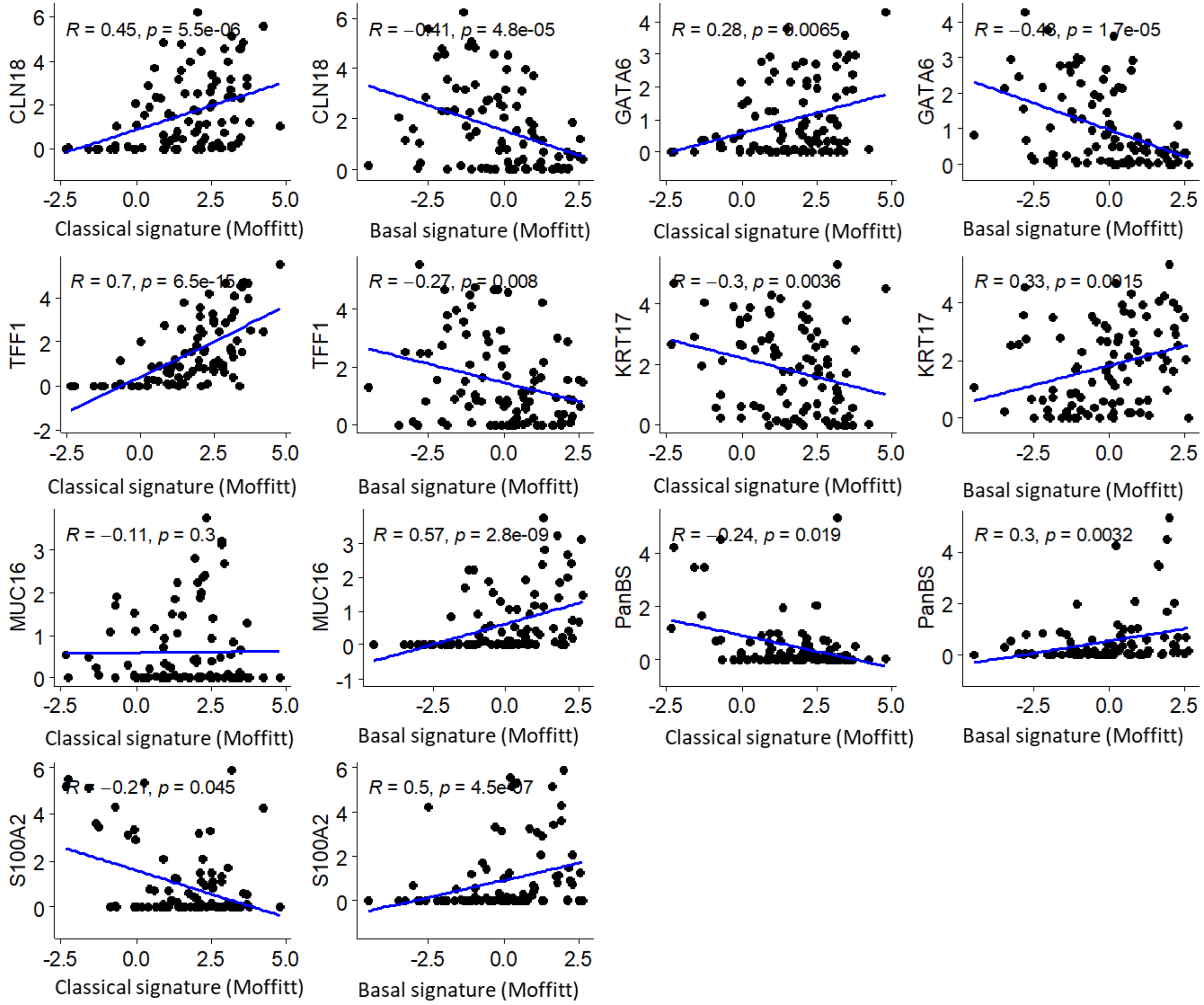

**Supplementary Figure 6.** K-means clustering of 44,024 tiles from 95 pancreatic ductal adenocarcinomas.

A

B

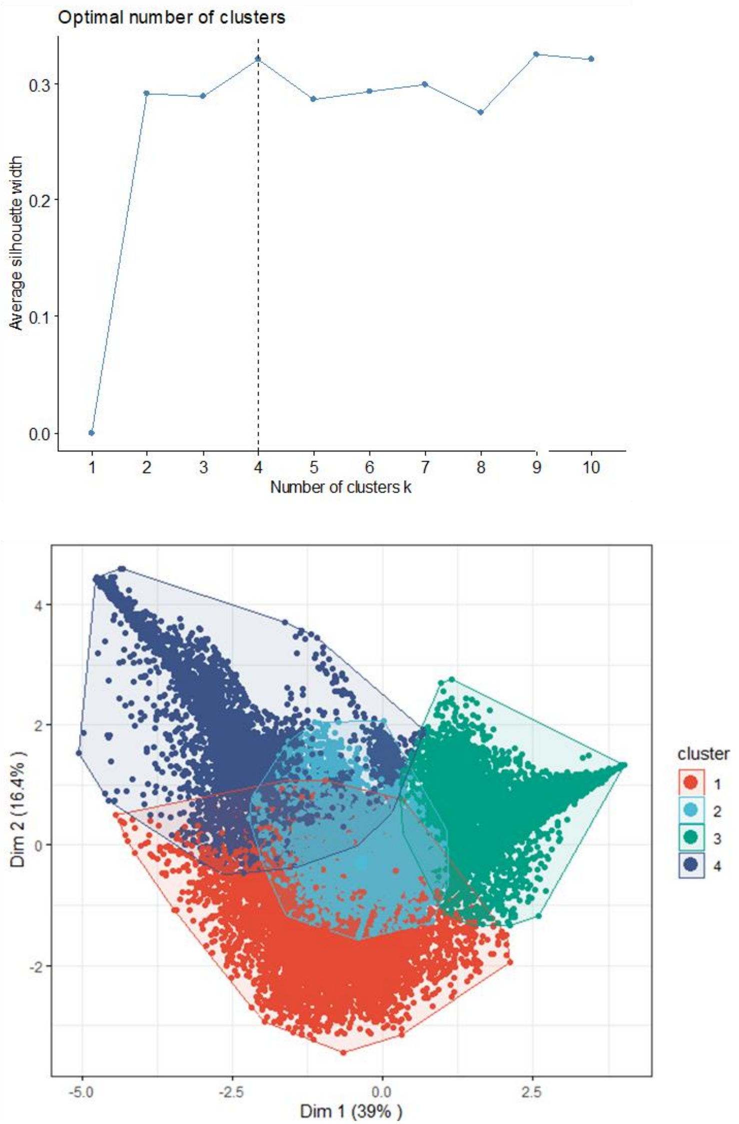

**Supplementary Figure 7.** Examples of representative areas of each cluster. A=Basal.MUC16, B=Basal.S100A2, C=Classical, D=Intermediate, E=Marker expression according to each cluster in representative areas.

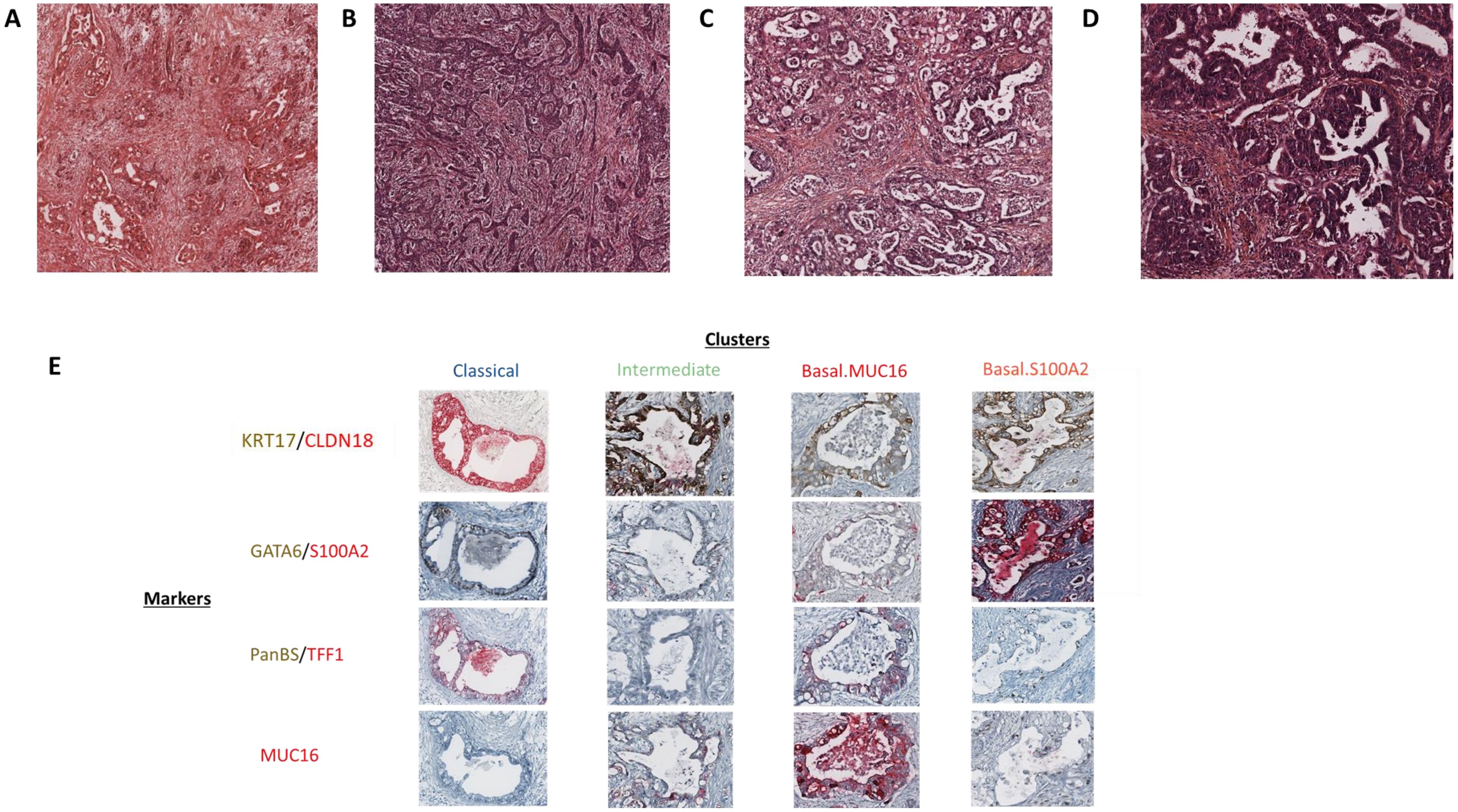

**Supplementary Figure 8.** Survival according to cluster. Multivariate analysis for the basal expression and clinicopathological factors regarding overall survival (n=95 patients) (A-B). Overall survival curves according to the predominant cluster in basal-expressing PDAC (n=61 patients) (C) and in basal-free PDAC (D).

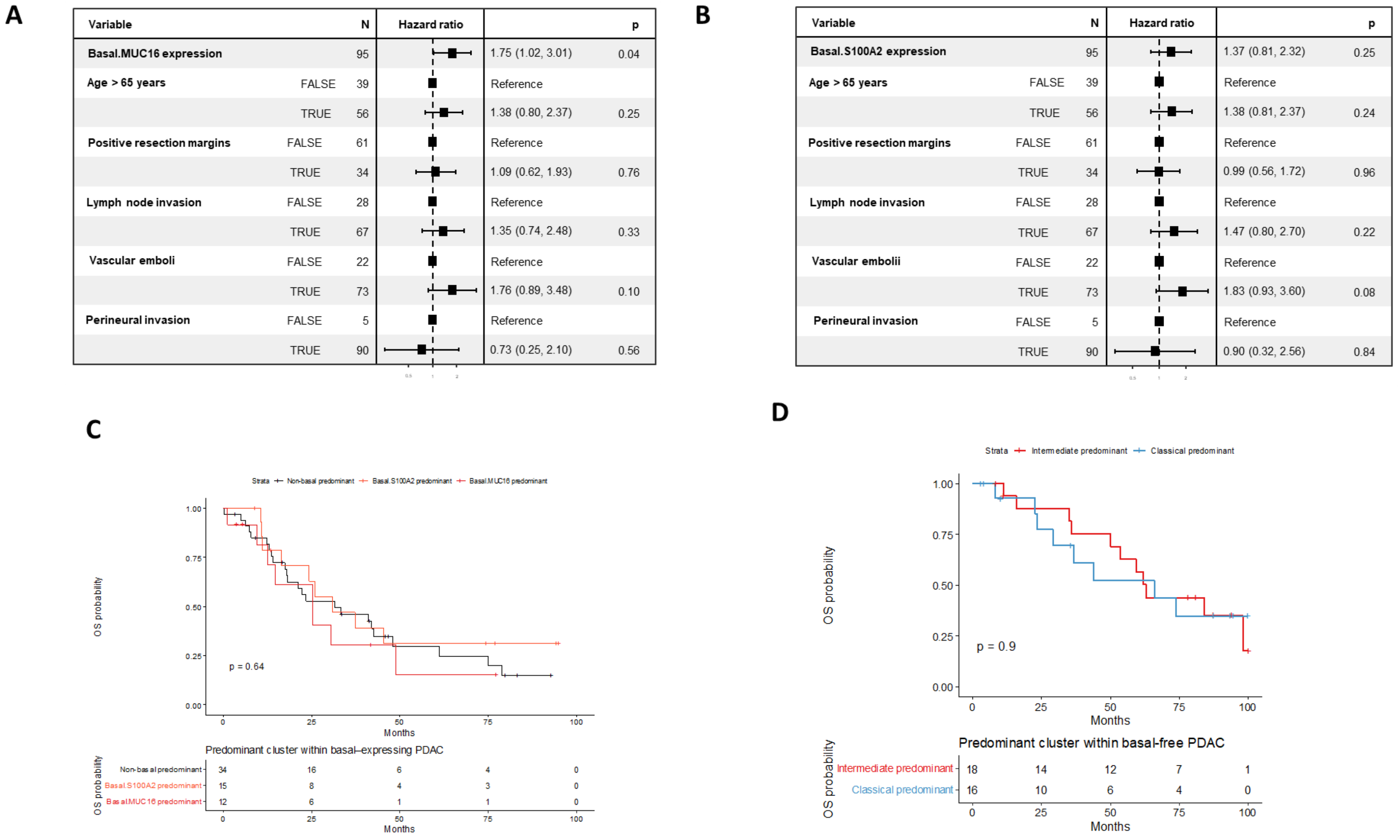

**Supplementary Figure 9.** Marker expression in pancreatic intraepithelial neoplasia.

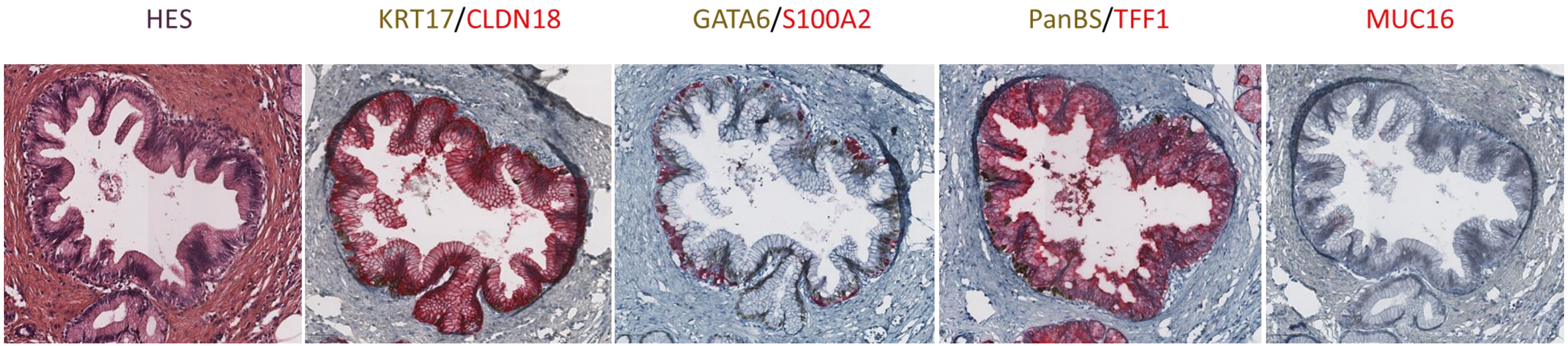

**Supplementary Figure 10.** Differential analysis between basal clusters. A. GSEA analysis showing the signaling pathways that are differentially regulated and statistically significant (adjusted p-value < 0.05) between Basal.MUC16 and Basal.S100A2 tumors B. Volcano plot showing the differentially expressed genes between early-onset and late-onset tumors. Dashed lines indicate the threshold of significant gene expression, defined as log2-transformed fold change ≤ −2.0 and ≥ 2.0 with adjusted p-value > 0.05.

A

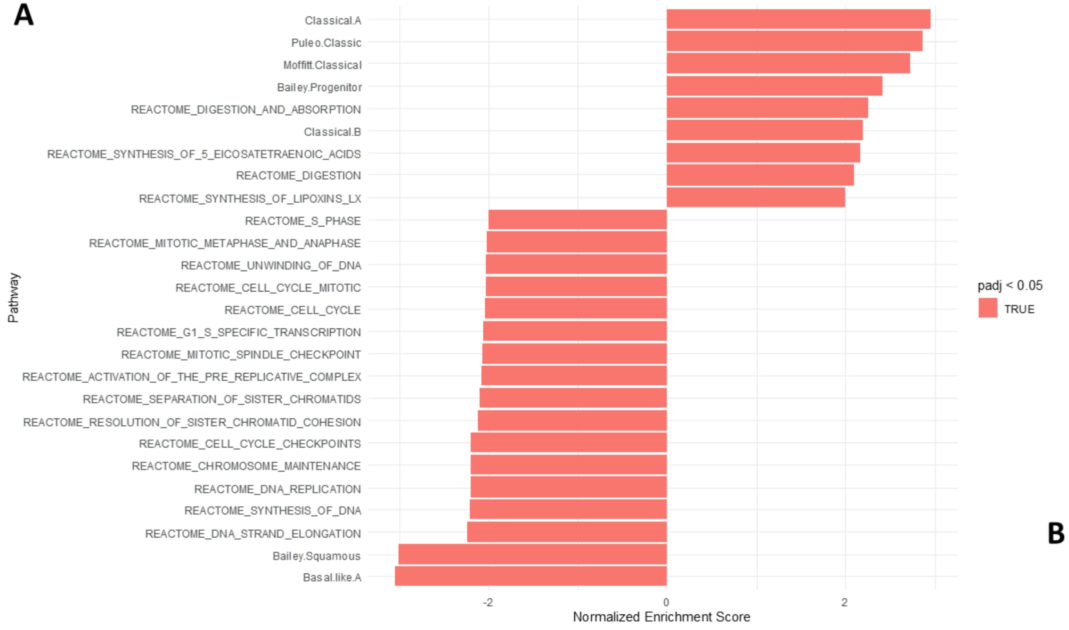


B

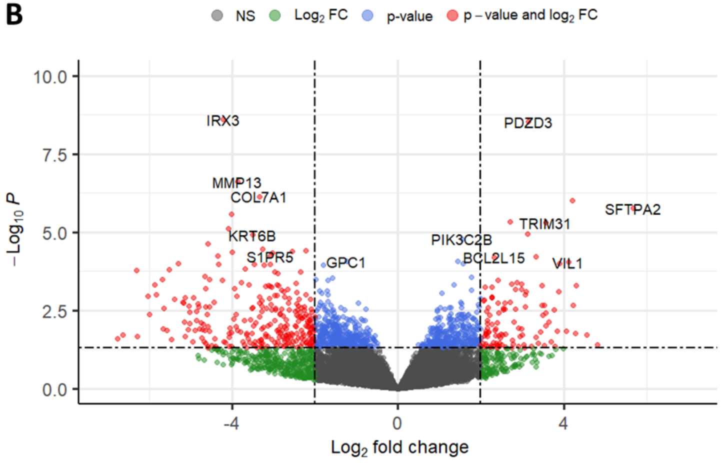
